# Childhood intelligence attenuates the association between biological ageing and health outcomes in later

**DOI:** 10.1101/588293

**Authors:** Anna J. Stevenson, Daniel L. McCartney, Robert F. Hillary, Paul Redmond, Adele M. Taylor, Qian Zhang, Allan F. McRae, Tara L. Spires-Jones, Andrew M. McIntosh, Ian J. Deary, Riccardo E. Marioni

## Abstract

The identification of biomarkers that discriminate individual ageing trajectories is a principal target in ageing research. Some of the most promising predictors of biological ageing have been developed using DNA methylation. One recent candidate, which tracks age-related phenotypes in addition to chronological age, is ‘DNAm PhenoAge’. Here, we performed a phenome-wide association analysis of this biomarker in a cohort of older adults to assess its relationship with a comprehensive set of both historical and contemporaneously-measured phenotypes. Higher than expected DNAm PhenoAge compared to chronological age, known as epigenetic age acceleration, was found to associate with a number of blood, cognitive, physical fitness and lifestyle variables, and with mortality. Notably, DNAm PhenoAge, assessed at age 70, was associated with cognitive ability at age 11, and with educational attainment. Adjusting for age 11 cognitive ability attenuated the majority of the cross-sectional later-life associations between DNAm PhenoAge and health outcomes. These results highlight the importance of early-life factors on healthy ageing.

## Introduction

A key objective in ageing research is the development of biomarkers that distinguish individuals on different ageing trajectories. Due to the distinct and calculable pattern of age-related changes in DNA methylation across the genome with chronological age, a number of DNA methylation-based biomarkers of ageing, or ‘epigenetic clocks’, have been developed. Accelerated epigenetic ageing has been linked with a number of age-related morbidities and with increased risk of mortality (1, 2). Whereas the first-generation of epigenetic clocks were developed using solely chronological age as the reference, a more recent effort additionally incorporated age-related phenotypes including blood cell profiles and inflammatory markers (3). This newer clock, termed DNAm PhenoAge, aimed to capture a truer and more efficacious epigenetic biomarker of physiological age, one which discriminates morbidity and mortality more definitively among individuals of the same chronological age.

DNAm PhenoAge was found to associate with diverse morbidities and mortality, with improved predictive power over other epigenetic clocks (3). However, many of the associations were with composite indices of health outcomes, rather than individual phenotypes. Moreover, the associations between DNAm PhenoAge and early life factors are currently unknown. It has been acknowledged that childhood and life-course traits and circumstances might have an enduring impact on later health. For example, greater childhood deprivation, lower childhood intelligence, relatively little formal education, and more manual adult occupations have been associated with increased morbidity and mortality in older age (2-4). Accordingly, for a more complete picture of the validity of DNAm PhenoAge, in addition to testing its relationship with individual ageing outcomes and mortality, it would be desirable to examine whether it can be predicted by these life history variables.

Here, we conduct a phenome-wide association study (PheWAS), in which multiple phenotypes are related to a single outcome, to investigate the link between accelerated DNAm PhenoAge and a comprehensive set of both historical and contemporaneously-assessed phenotypes in a large, longitudinal cohort study of ageing: the Lothian Birth Cohort 1936 (LBC1936). This cohort is unusually valuable because data are available on their general cognitive ability and social circumstances at age 11 which can inform the understanding of possible early life confounders of later-life outcomes.

## Methods

### Study population

The Lothian Birth Cohort 1936 (LBC1936) is a longitudinal study of ageing. The cohort comprises a community-dwelling sample of participants born in 1936, most of whom undertook a general intelligence test - the Moray House Test No. 12 - in 1947, aged around 11 years. In total 1,091 participants were recruited to the study at a mean age of about 70 years, and have subsequently been re-examined at three furthers waves, aged around 73, 76 and 79 years. Participants have been comprehensively phenotyped at each wave of the study with data collected on cognitive measures, physical and health outcomes, genetics, lifestyle factors and psycho-social aspects of ageing. Full details on the background, recruitment and data collection procedures of the study are provided elsewhere (4, 5).

### Ethics and consent

Ethical permission for LBC1936 was obtained from the Multi-Centre Research Ethics Committee for Scotland (MREC/01/0/56) and the Lothian Research Ethics Committee (Wave 1: LREC/2003/2/29) and the Scotland A Research Ethics Committee (Waves 2, 3 and 4: 07/MRE00/58). Written informed consent was obtained from all participants.

### LBC1936 DNA methylation

The LBC1936 methylation profiling has been fully detailed previously (6, 7). Briefly, DNA was extracted from whole-blood samples at Wave 1 of the study and methylation was measured at 485,512 probes. Quality control analysis resulted in the removal of CpG sites with a low detection rate (<95% at p<0.01). Probes with low quality (inadequate hybridisation, bisulfite conversion, nucleotide extension and staining signal) were additionally identified and removed after manual inspection of the array control probe signals. Finally, probes with a low call rate (<450,000 probes detected at p<0.01), XY probes, and samples in which the predicted sex did not match the reported sex, were excluded.

### DNAm PhenoAge

The DNAm PhenoAge biomarker was developed in a two-step process by Levine et al (3). Briefly, a novel measure of ‘phenotypic age’ was developed using penalised regression where the hazard of mortality was regressed on 42 clinical markers from the third National Health and Nutrition Examination Survey (NHANES-III). The optimal model selected nine variables (albumin, creatinine, serum glucose (HbA_1c_), C-reactive protein (CRP), percentage lymphocytes, mean cell volume, red cell distribution width, alkaline phosphatase and white blood cell count) in addition to chronological age for inclusion in the phenotypic age predictor. An epigenetic biomarker of phenotypic age was then developed using the Invecchiare in Chianti (InCHIANTI) cohort (8), by regressing computed phenotypic age on whole-blood DNA methylation data, producing an estimate of DNAm PhenoAge based on methylation profiles at 513 CpGs.

DNAm PhenoAge was calculated in LBC1936 by multiplying methylation beta values with the regression weights from the above analysis (3). One CpG (cg06533629) from the 513 used in the original computation was not available in the LBC1936 methylation data. At Wave 1, 889 individuals within the LBC1936 cohort had full methylation data available for the calculation of DNAm PhenoAge.

### Phenotypic data

The PheWAS included 107 phenotypes broadly associated with health and wellbeing. The phenotypes encompassed seven subgroups: blood, cardiovascular, cognitive, personality and mood, lifestyle, physical, and life-history, and were measured on a binary (n=15), continuous (n=89) or ordinal (n=3) scale. Six of the phenotypes included in the PheWAS (white cell counts, blood glucose, CRP, creatinine, albumin and mean cell volume) were incorporated in the original phenotypic age predictor.

Descriptive statistics for the phenotypes are presented in **Supplementary file 1**. Data collection protocols are detailed in **Supplementary file 2** and have been described fully previously (9).

### Statistical Analysis

DNAm PhenoAge acceleration (DNAm PhenoAgeAccel) – defined as the residuals resulting from regressing DNAm PhenoAge on chronological age - was calculated for all participants at LBC1936’s Wave 1, at a mean age of 70 years.

Linear regression models were used to obtain the associations between the continuous variables with DNAm PhenoAgeAccel. All continuous variables were scaled to have a mean of zero and unit variance to ensure comparable effect sizes across all traits. Generalised linear models with a logit link function (logistic regression) were used to investigate the association between the binary variables and DNAm PhenoAgeAccel, and ordinal regression models were used for the ordered categorical measures of smoking (3 levels), physical activity (5 levels) and occupational social class (6 levels). DNAm PhenoAgeAccel was the independent variable of interest in each regression model. Height and smoking status (Wave 1) were included as covariates in the models for lung function (forced expiratory volume FEV_1_; forced vital capacity: FVC; forced expiratory ratio: FER; and peak expiratory flow: PEF). All models were adjusted for chronological age and sex. To investigate the influence of childhood cognitive ability, all models that showed significant associations with DNAm PhenoAgeAccel were repeated adjusting for age-11 IQ scores.

In the longitudinal analysis, linear mixed-effects models were used to assess if baseline DNAm PhenoAgeAccel was associated with longitudinal change over the four waves of data (∼70 years to ∼79 years) in a subset of the cognitive and physical phenotypes that are known to decline with age and correlate with functional impairment. Here, Wave 1 DNAm PhenoAgeAccel was included as a fixed-effect interaction with chronological age, and participant was added as a random-effect intercept term. As above, height and smoking status were included in the models for lung function, and all models co-varied for sex. Cox proportional-hazards model was implemented to analyse the effect of DNAm PhenoAgeAccel on survival (time-to-death).

Correction for multiple testing was applied using the false discovery rate (FDR) method for each group of variables (10).

Statistical analysis was conducted in R version 3.5.0 using the ‘lm’ and ‘glm’ function in the ‘stats’ library and the ‘lme4’, ‘lmerTest’, ‘rms’ and ‘Survival’ packages (11-16).

## Results

### Cohort information

Details of the baseline (Wave 1) characteristics of LBC1936 are presented in **Supplementary file 1**. 49.8% of the cohort was female. Mean chronological age for both males and females was 69.5 years (SD 0.8) and mean DNAm PhenoAge was 57.8 years (females = 56.7 [SD = 8.1], males = 58.8 [SD 8.2]). The discrepancy between the chronological and epigenetic age measures is probably reflective of the overall good health of the cohort.

### PheWAS

Only associations with an FDR-corrected significant p-value (<0.05) are presented here and in **Figure 1**. Full results are presented in **Supplementary file 3** and **Supplementary file 4.**

**Figure 1.**
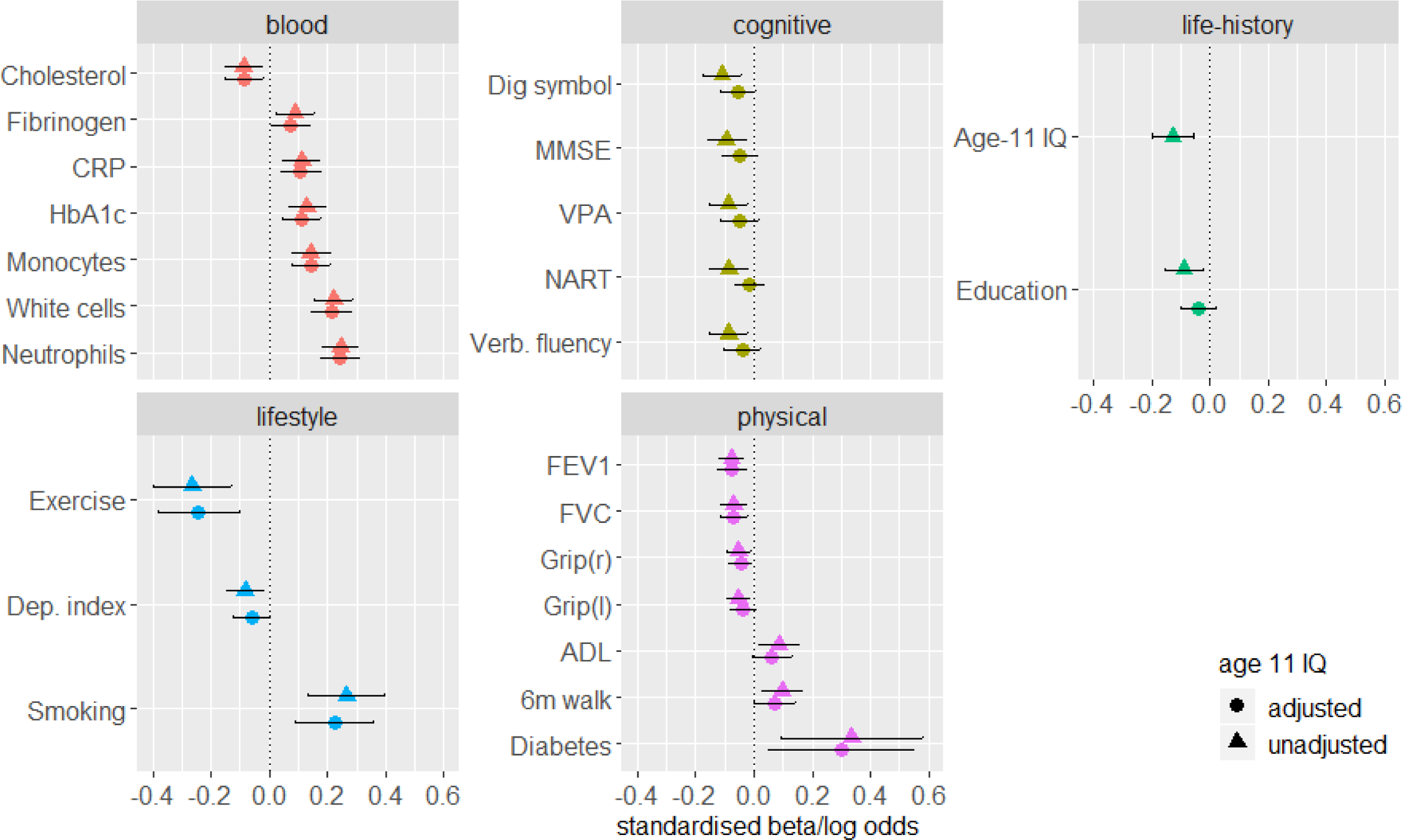
FDR-corrected significant associations between DNAm PhenoAgeAccel and blood, cognitive, lifestyle, physical and life-history variables. Standardised model β coefficients (for continuous variables) or log odds (for binary variables) are presented along the x-axes. Phenotypes are presented along the y-axis. Error bars show the 95% confidence interval. CRP: C-reactive protein; Dep: deprivation; VPA: verbal paired associates; verb: verbal; NART: National Adult Reading Test; MMSE: Mini-Mental State Examination; grip: grip strength; FEV1: forced expiratory volume in 1 second; FVC: forced vital capacity; ADL: activities of daily living (Townsend Disability Scale).

#### Blood

Significant associations between DNAm PhenoAgeAccel and blood phenotypes are presented in **Figure 1**. Full results are presented in **Supplementary file 3: Figure 1** and **Supplementary file 4**. Of the six measures included in the phenotypic age reference, three significant positive associations were found with DNAm PhenoAgeAccel: white cell counts (β = 0.22, SE = 0.03, p = 2.5×10^−9^) HbA_1c_ (β = 0.13, SE = 0.03, p = 5.8×10^−4^) and C-reactive protein (CRP; β = 0.11, SE = 0.03, p = 0.006).

Significant positive relationships were additionally identified between DNAm PhenoAgeAccel and neutrophils (β = 0.25, SE = 0.03, p = 2.9×10^−12^), monocytes (β = 0.14, SE = 0.03, p = 1.6×10^−4^) and fibrinogen (β = 0.09, SE = 0.03, p = 0.043). A negative association was found between DNAm PhenoAgeAccel and total cholesterol levels (β = −0.09, SE = 0.03, p = 0.043).

#### Cardiovascular

No significant associations were found between DNAm PhenoAgeAccel and any of the cardiovascular variables (**Supplementary file 3: Figure 2**, FDR-corrected p ≥ 0.12).

**Figure 2.**
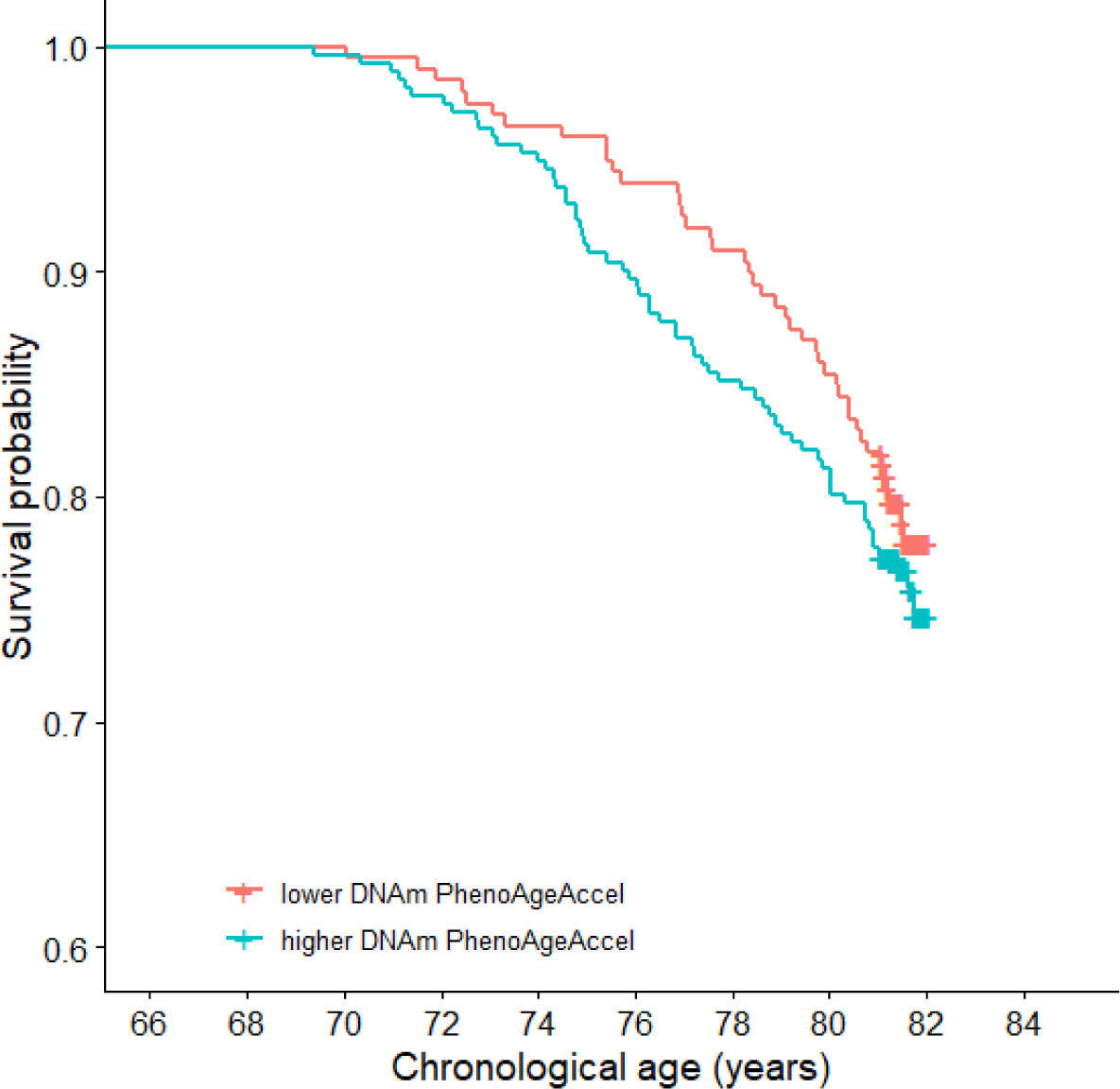
Survival probability by quartiles of DNAm PhenoAgeAccel adjusted for sex and chronological age.

#### Cognitive

Significant associations between DNAm PhenoAgeAccel and cognitive phenotypes are presented in **Figure 1**. Full results are presented in **Supplementary file 3: Figure 3** and **Supplementary file 4**. Higher DNAm PhenoAgeAccel associated with lower scores on one test of the processing speed domain (digit symbol coding: β = −0.11, SE = 0.03, p = 0.015), one test of the memory domain (verbal paired associates: β = −0.09, SE = 0.03, p = 0.034), the mini-mental state examination (MMSE; β = − 0.10, SE = 0.03, p = 0.034) and two tests of crystallised ability (national adult reading test [NART]: β = −0.09, SE = 0.03, p = 0.034; verbal fluency: β = −0.09, SE = 0.03, p = 0.034).

#### Personality and mood

No significant associations were found between DNAm PhenoAgeAccel and any of the personality and mood phenotypes **(Supplementary file 3: Figure 4**).

#### Physical

Significant associations between DNAm PhenoAgeAccel and physical phenotypes are presented in **Figure 1**. Full results are presented in **Supplementary file 3: Figure 5** and **Supplementary file 4**. We found significant inverse associations between DNAm PhenoAgeAccel and FEV_1_ (β = −0.07, SE = 0.02, p = 0.023), FVC (β = −0.07, SE = 0.02, p = 0.023), and grip strength in both right and left hands (both: β = −0.05, SE = 0.02, p = 0.045). Higher DNAm PhenoAgeAccel was associated with a diagnosis of diabetes (OR = 1.39, 95% CI [1.01, 1.78], p = 0.038), a slower six metre walk time (β = 0.01, SE = 0.03, p = 0.038), and a higher score on the Townsend’s Disability Scale (activities of daily living; β = 0.09, SE = 0.03, p = 0.045).

#### Lifestyle

Significant associations between DNAm PhenoAgeAccel and lifestyle phenotypes are presented in **Figure 1**. Full results are presented in **Supplementary file 3: Figure 6** and **Supplementary file 4**. A significant inverse association was found between baseline DNAm PhenoAgeAccel and deprivation index (β = −0.08, SE = 0.03, p = 0.025). Additionally, a higher DNAm PhenoAgeAccel was associated with lower levels of physical activity (OR = 0.77, 95% CI [0.67, 0.88], p = 0.0003) and with higher odds of being either a current or an ex-smoker, compared to a never smoker (OR = 1.31, 95% CI [1.15, 1.49], p = 0.0003).

#### Life-history

Significant associations between DNAm PhenoAgeAccel and life-history phenotypes are presented in **Figure 1**. Full results are presented in **Supplementary file 3: Figure 7** and **Supplementary file 4**. A higher baseline DNAm PhenoAgeAccel was associated with a lower age-11 IQ (β = −0.13, SE = 0.03, p = 0.001) and fewer years of education (β = −0.09, SE = 0.03, p = 0.013).

### Longitudinal association between DNAm PhenoAgeAccel and phenotypes

All the cognitive and physical fitness measures included in the longitudinal analysis showed changes over time that were consistent with declining health (**Supplementary file 5**). The rate of decline ranged from 0.02 SDs per year (digit span backwards) to 0.08 SDs per year (telomere length). Six metre walk time increased by 0.1 SDs per year (all p ≤ 2×10^−8^).

Baseline DNAm PhenoAgeAccel was not found to associate with subsequent change in any of the assessed phenotypes (FDR-corrected p ≥ 0.322, **Table 1**).

**Table 1.** Longitudinal associations between baseline DNAm PhenoAgeAccel and phenotypes.

| Phenotype | Standardised $\beta$ | Standard error | Raw p | FDR-corrected p |
| --- | --- | --- | --- | --- |
| *Grip strength (r)* | 0.010 | 0.008 | 0.189 | 0.448 |
| *Grip strength (l)* | 0.008 | 0.008 | 0.288 | 0.448 |
| *Forced expiratory volume (1s)* | 0.009 | 0.008 | 0.261 | 0.448 |
| *Forced vital capacity* | 0.011 | 0.009 | 0.192 | 0.448 |
| *Forced expiratory ratio* | 0.021 | 0.015 | 0.175 | 0.448 |
| *Peak expiratory flow* | 0.006 | 0.011 | 0.613 | 0.859 |
| *Digit span backwards* | <0.001 | 0.013 | 0.952 | 0.952 |
| *Symbol search* | -0.003 | 0.013 | 0.821 | 0.952 |
| *Digit symbol coding* | 0.019 | 0.009 | **0.046** | 0.322 |
| *Matrix reasoning* | 0.004 | 0.013 | 0.713 | 0.908 |
| *Letter number sequencing* | -0.018 | 0.013 | 0.180 | 0.448 |
| *Block design* | -0.011 | 0.011 | 0.282 | 0.448 |
| *6m walk time (s)* | <0.001 | 0.014 | 0.945 | 0.952 |
| *Telomere length* | -0.025 | 0.012 | **0.041** | 0.322 |

### DNAm PhenoAge and survival

We tested the association of DNAm PhenoAge with all-cause mortality (n_deaths_= 209, over 9 years of follow-up, average age of death = 76.8 years, SD 3.3) and found that a higher DNAm PhenoAgeAccel was significantly associated with risk of death (HR = 1.17 per SD increase in DNAm PhenoAgeAccel, 95% CI [1.02, 1.34], p = 0.025). A Kaplan-Meier survival curve for DNAm PhenoAgeAccel, split into highest and lowest quartiles, is presented in **Figure 2**, illustrating the higher mortality risk for those with a higher DNAm PhenoAgeAccel.

### Adjusting for Age-11 IQ

To test for potential confounding of the associations by childhood intelligence, the models for all of the significant associations identified in the PheWAS were re-run adjusting for age-11 IQ. Results are presented in **Figure 1** and **Table 2.**

**Table 2.**
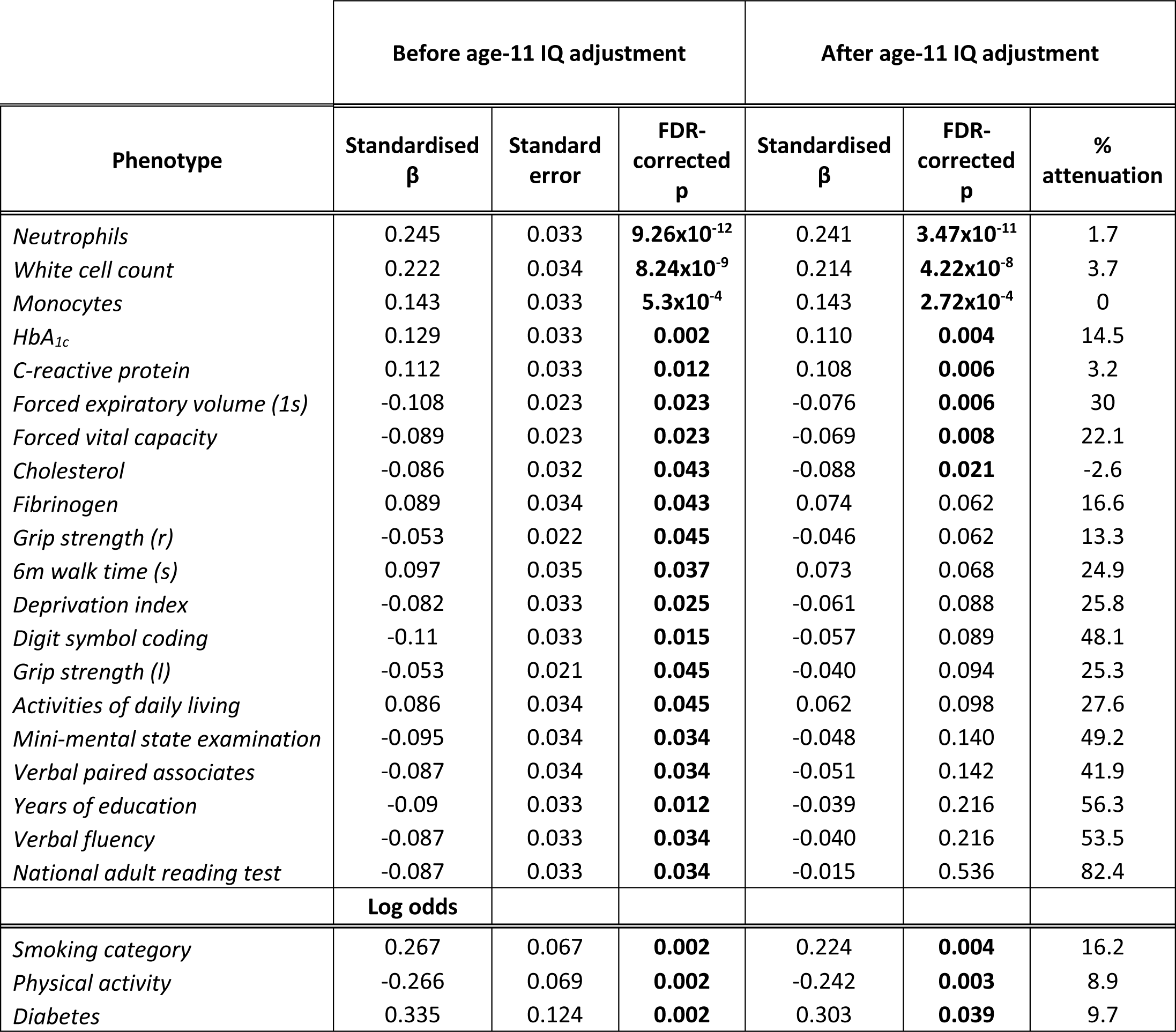
Results before and after adjusting models for age-11 IQ. Standardised β are presented for continuous variables and log odds for binary or ordinal phenotypes. FDR-corrected significant results are highlighted in bold.

The associations with 12 out of the 23 phenotypes that were originally found to be significant in the PheWAS, became non-significant upon adjustment, inclusive all of the cognitive associations. Of the associations that became non-significant, the effect sizes were attenuated by a mean of 39% (range: 13.4% to 82.4%). The survival model was also no longer significant following adjustment for age-11 IQ (HR = 1.13, 95% CI [0.98, 1.31], p = 0.08).

All of the associations with blood phenotypes, excepting fibrinogen, were still significant following adjustment for age-11 IQ, as were the lung function measures, smoking status, physical activity and diabetes.

## Discussion

In this study we performed a comprehensive PheWAS to investigate the associations between 107 phenotypes with a new epigenetic predictor - DNAm PhenoAge - in a large cohort of older adults. We identified significant correlations at a mean age of 70 years between accelerated DNAm PhenoAge and a number of blood-based, physical, cognitive, and lifestyle phenotypes, in addition to mortality. Importantly, we found that the life-history variables of general cognitive ability, measured at age 11, and number of years of education, related to DNAm PhenoAge at age 70. Moreover, adjustment for age 11 cognitive ability attenuated the majority of the cross-sectional later-life associations between DNAm PhenoAge and health outcomes. Most of the blood-based phenotypes – many of which were used to derive the PhenoAge predictor – remained significant, along with some measures of physical fitness and smoking status. We found no link between baseline DNAm PhenoAgeAccel and longitudinal change in any of the assessed phenotypes.

### PheWAS results

#### Blood

Higher DNAm PhenoAgeAccel associated with raised levels of various blood-based immune and inflammatory cells (white cell counts, monocytes and neutrophils), alongside elevated blood glucose (HbA_1c_), and two inflammatory mediators - fibrinogen and CRP. The associations identified with white cell counts, glucose and CRP are largely unsurprising given the derivation of phenotypic age. Conversely, an inverse relationship was found with total cholesterol. High total cholesterol concentrations have previously been shown to have protective effects in older age, associating with longevity (17). Taken together, these results suggest that DNAm PhenoAge captures features of age-related inflammation and immunosenescence. These findings seem congruous with the ‘inflamm-aging’ theory, which posits that chronic, low-grade, sterile inflammation is a central driver of the ageing process (18).

#### Cognitive

As cognitive decline is one of the most feared aspects of growing older, characterising its association with a predictor of ageing outcomes is important. The negative association between DNAm PhenoAgeAccel and the MMSE suggests a potential link between a higher epigenetic age and lower cognitive ability in older age; however only two out of eleven of the tests of cognitive function domains that typically decline in older age (visual paired associates [memory] and digit-symbol coding [processing speed]) were found to have a significant inverse relationship with DNAm PheonAgeAccel. We found that higher DNAm PhenoAgeAccel was associated with lower scores on two cognitive tests that comprise measures of crystallised intelligence (NART and verbal fluency). Given that crystallised ability typically remains relatively stable throughout the life course (19), this result seems reflective of a link between accelerated biological ageing and generalised lower scores in the crystallised domain.

#### Physical

Akin to the findings from Levine et al. that higher DNAm PhenoAge is associated with increased physical functioning problems, we found significant associations between more impaired scores on measures of age-related physical fitness – gait speed, grip strength, lung function and the Townsend Disability Scale (activities of daily living) - with higher DNAm PhenoAgeAccel. The only overt disease phenotype found to correlate with DNAm PhenoAgeAccel was diabetes. This association may be a result of the inclusion of HbA_1c_ in the phenotypic age measure capturing recognised alterations in blood glucose seen in diabetes.

#### Lifestyle

We observed that lower levels of physical activity and ranking on a more deprived socioeconomic scale correlated with an increased DNAm PhenoAgeAccel. The association with social deprivation is conceivably due to its link with more biologically direct risk factors for ageing and morbidity. For instance, various lifestyle variables have been shown to associate with socioeconomic status, including smoking and BMI – two of the largest risk factors for a host of age-related diseases (20, 21). The association identified between smoking status and DNAm PhenoAgeAccel is not surprising given the significant impact smoking has on disease risk and life expectancy and on DNA methylation (22). Overall, these findings iterate the protective effects of a higher socioeconomic status, exercise and abstention from smoking in the mitigation of susceptibility to age-related decline.

#### Life-history

Of the assessed life-history variables, fewer education years correlated with an increased DNAm PhenoAgeAccel as did lower age-11 IQ scores. While these may be independent associations, childhood cognitive function is a substantial predictor of educational attainment (23). The association between DNAm PhenoAgeAccel with IQ measured almost 60 years previously is a key finding and is indicative of a lifelong, enduring association between cognition and DNAm PhenoAgeAccel rather than one specific to ageing. This bolsters cognitive epidemiology findings indicating that general intelligence in childhood, as measured by psychometric tests, predicts, or is associated with, substantial life-course differences in health and morbidity (24, 25). Various, non-exclusive, mechanisms are thought to govern this association, including better health literacy and disease management, higher socioeconomic standing, and the ‘system integrity’ hypothesis which postulates that higher scores on cognitive ability tests are capturing a systemic level of good functioning rather than isolated brain efficiency (26). It is possible that individual differences in DNAm PhenoAgeAccel in older age are, in part, caused by intelligence differences over the lifecourse, or that both are a result of a shared genetic architecture or early environmental event.

### Longitudinal association and mortality

Similar to Levine et al. original paper, we found accelerated DNAm PhenoAge to be associated with a higher risk of mortality. However, we found no evidence to suggest that DNAm PhenoAgeAccel associates with longitudinal changes in ageing phenotypes, limiting its potential as a prospective biomarker of ageing.

### Adjusting for age-11 IQ

All of the associations found between DNAm PhenoAgeAccel with concurrent cognitive measures ceased to be significant following adjustment for age-11 IQ. Comparably, all of the associations with the physical fitness phenotypes, excepting the lung function measures, were attenuated with P>0.05. This suggests that the relationship found between DNAm PhenoAgeAccel and contemporaneous cognition in older age, in addition to walking speed, grip strength and physical ability, is partially mediated through childhood cognitive ability, solidifying the aforementioned lifetime association between intelligence and health. The associations with education duration and socioeconomic status were also attenuated and no longer significant upon adjustment. As previously mentioned, childhood IQ has been shown to predict educational outcomes and the two are significantly genetically correlated (27, 28). Additionally, it has been demonstrated that lower childhood IQ associates with adult socioeconomic deprivation and reciprocally, higher IQ in childhood was found to be a predictor of upward social mobility (29-31). Childhood IQ has frequently been associated with both all-cause (32) and cause-specific mortality (33-36) and the survival model in our analyses also became non-significant following adjustment. These findings are consistent with the theory of confounding or reverse causation; that is, cognitive ability at age 11 accounts for most of the cross-sectional associations between epigenetic age acceleration and various health and wellbeing phenotypes in later-life.

Though the majority of the formerly significant relationships were attenuated and became non-significant in the adjusted models, the associations with most of the blood phenotypes, including the three that were incorporated in the phenotypic age predictor, remained significant. Additionally, though the effect sizes attenuated by 30% and 22% respectively, the associations with the lung function measures of FEV_1_ and FVC remained significant. There was little attenuation of the associations with levels of physical activity and not smoking, perhaps exhibitive of their importance in terms of biological ageing.

### Strengths and Limitations

This is the first independent test of DNAm PhenoAge in a large cohort of older adults with the availability of historical variables, as well as longitudinal measures for an extensive number of health and ageing-related phenotypes. Moreover, these data are available across the 8th decade, a time when risk of dementia and functional decline increases substantially. Critically, the availability of childhood IQ measures enabled us to show that many cross-sectional associations between DNAm PhenoAge and health are confounded by early life cognitive ability.

LBC1936 are a predominantly healthy older ageing cohort, reflected by the young estimation of DNAm PhenoAge compared to chronological age, which might preclude the generalisation of these findings to the broader ageing population in which manifold co-existing morbidities are commonly prevalent. Furthermore, most of the disease assessments within the study are concluded from self-reports which are often unreliable, limiting their use as indicators of verifiable pathologies. These aspects perhaps hindered additional findings of disease-related associations, and future studies could consider DNAm PhenoAge associations in larger, longitudinal cohorts with clinical end-points.

### Conclusion

We have verified associations between an innovative and novel marker of epigenetic age and a number of pertinent, proxy health-related phenotypes and mortality in older adults. Additionally, intelligence at age 11 and educational attainment were found to associate with DNAm PhenoAge at age 70. Most notably, adjustment for childhood cognitive ability attenuated over half of the late-life associations of health and DNAm PhenoAge by 13%-82%. While it does seem DNAm PhenoAge independently captures some measures of age-related functional fitness and blood-based phenotypes, future studies utilising it as a biomarker should be aware of potentially mediating factors in the interpretation of results.

## Acknowledgements

The authors thank all LBC1936 study participants and research team members who have contributed, and continue to contribute, to ongoing LBC1936 studies.

