## Supplementary file 1 for "Childhood intelligence attenuates the association between biological ageing and health outcomes in later"

**Supplementary file 1.** Details of the Lothian Birth Cohort 1936 phenotypes at Wave 1 (~70 years).

*Abbreviations*: SD: standard deviation; WMS: Wechsler Memory Scale; WAIS: Wechsler Adult Intelligence Scale; IPIP: International Personality Item Pool; NEO-FFI: Neuroticism-Extraversion-Openness Five-Factor Inventory; SIMD: Scottish Index of Multiple Deprivation.

| Phenotype | n | | % |
|---|---|---|---|
| Sex (female) | 543 | | 49.8 |
| | n | Mean | SD |
| Age (years) | 1091 | 69.5 | 0.83 |
| DNAmPhenoAge (years) | 889 | 57.8 | 8.2 |
| **BLOOD** | | | |
| Haemoglobin 130-180 g/L | 1064 | 145.2 | 13 |
| Red cell count 4.5-6.5 10^12/L | 1063 | 4.61 | 0.41 |
| Haematocrit 0.40-0.54 ratio | 1063 | 0.42 | 0.04 |
| Mean cell volume 78.0-98.0 fL | 1063 | 91.12 | 4.45 |
| White cell count 4.0-11.0 10^9/L | 1063 | 7.04 | 2.26 |
| Neutrophil count 2.0-7.5 10^9/L | 1063 | 4.44 | 1.56 |
| Lymphocyte count 1.5-4.0 10^9/L | 1063 | 1.88 | 1.33 |
| Monocyte count 0.2-0.8 10^9/L | 1063 | 0.53 | 0.19 |
| Eosinophil count 0.04-0.4 10^9/L | 1063 | 0.17 | 0.13 |
| Basophil count 0.01-0.1 10^9/L | 1063 | 0.04 | 0.03 |
| Platelet count 150-350 10^9/L | 1062 | 274.18 | 65.9 |
| Prothrombin time 8.0-11.0 secs | 1052 | 10.16 | 2.78 |
| Prothrombin time ratio 0.8-1.2 ratio | 1052 | 1 | 0.28 |
| Activated partial thromboplastin time 27.0-38.0 secs | 1052 | 28.59 | 3.07 |
| Activated partial thromboplastin time ratio 0.8-1.2 ratio | 1052 | 0.94 | 0.11 |
| Fibrinogen 1.5-4.0 g/L | 1052 | 3.28 | 0.64 |
| Vitamin B12 200-900 ng/L | 910 | 414.67 | 140.76 |
| Serum folate 5-20 ug/L | 912 | 12.82 | 6.29 |
| Red cell folate 257-800 ug/l | 1049 | 412.68 | 124.15 |
| Urea 2.5-6.6 mmol/L | 1062 | 6.03 | 1.59 |
| Creatinine 60-120 umol/L | 1062 | 77.85 | 17.17 |
| Sodium 135-145 mmol/L | 1061 | 140.83 | 2.62 |
| Potassium 3.6-5 mmol/L | 821 | 4.38 | 0.36 |
| Albumin 35-50 g/L | 1059 | 44.67 | 3.1 |
| Calcium 2.1-2.6 mmol/L | 1057 | 2.35 | 0.09 |
| Triglyceride (0.8 - 2.1 mmol/L) | 965 | 1.64 | 0.78 |
| Cholesterol (mmol/L) | 1055 | 5.45 | 1.15 |
| High-density lipoprotein cholesterol 0.9-1.4 mmol/L | 970 | 1.52 | 0.44 |
| Cholesterol: HDLC Ratio | 966 | 3.76 | 1.07 |
| $HbA_{1C}$ 5.0-6.5 %total | 1062 | 5.93 | 0.73 |
| C-reactive protein 0-10 mg/L | 1054 | 5.26 | 6.68 |
| Thyroid stimulating hormone 0.5-4.7 mU/L | 1061 | 2.09 | 1.65 |
| Free thyroxine 9-24 pmol/L | 1059 | 15.37 | 2.49 |

| | | | |
|---|---|---|---|
| Total triiodothyronine 1.0-2.6 nmol/L | 943 | 2.14 | 0.38 |
| **CARDIOVASCULAR** | | | |
| High blood pressure (yes/no) | 433/658 | - | - |
| High cholesterol (yes/no) | 386/704 | - | - |
| Cardiovascular disease history (yes/no) | 268/823 | - | - |
| Problems with blood circulation (yes/no) | 156/932 | - | - |
| History of stroke (yes/no) | 54/1037 | - | - |
| Family history of heart disease, stroke or problems with blood vessels (yes/no) | 672/413 | - | - |
| Mean diastolic blood pressure - sitting | 1088 | 81 | 10.31 |
| Mean diastolic blood pressure - standing | 1086 | 85 | 10.25 |
| Mean systolic blood pressure - sitting | 1088 | 150 | 19.2 |
| Mean systolic blood pressure - standing | 1086 | 148 | 19.54 |
| **COGNITIVE** | | | |
| Mini-Mental State Examination (MMSE) total score | 1090 | 28.79 | 1.43 |
| WMS III - Logical memory (I + II) total score | 1087 | 71.46 | 17.96 |
| WMS III - Spatial span total | 1084 | 14.72 | 2.83 |
| WMS III - Verbal paired associates total score (I + II) | 1050 | 26.44 | 9.13 |
| WAIS III - Symbol search | 1086 | 24.75 | 6.29 |
| WAIS III - Digit-symbol coding total score | 1086 | 56.59 | 12.93 |
| Simple reaction time mean | 1085 | 0.28 | 0.057 |
| Log10 transformation simple reaction time mean | 1085 | -0.57 | 0.078 |
| Four choice reaction time mean | 1084 | 0.64 | 0.086 |
| Inspection time total correct responses | 1041 | 112.14 | 10.99 |
| WAIS III - Matrix reasoning total score | 1086 | 13.49 | 5.13 |
| Verbal fluency total score | 1087 | 42.42 | 12.54 |
| WAIS III - Letter-number sequencing | 1079 | 10.92 | 3.16 |
| WAIS III - Digit span backwards | 1090 | 7.73 | 2.26 |
| National Adult Reading Test (number correct) | 1089 | 34.48 | 8.15 |
| Wechsler Test of Adult Reading (number correct) | 1089 | 41.02 | 7.17 |
| WAIS III - Block design total score | 1085 | 33.79 | 10.32 |
| **PERSONALITY AND MOOD** | | | |
| Hospital Anxiety and Depression Scale - Anxiety score | 1089 | 4.89 | 3.18 |
| Hospital Anxiety and Depression Scale - Depression score | 1086 | 2.8 | 2.23 |
| Hospital Anxiety and Depression Scale - Total score | 1086 | 7.68 | 4.52 |
| IPIP Extraversion total score | 954 | 21.31 | 7.07 |
| IPIP Agreeableness total score | 952 | 31.08 | 5.41 |
| IPIP Conscientiousness total score | 952 | 28.23 | 5.99 |
| IPIP Emotional stability total score | 950 | 24.59 | 7.66 |
| IPIP Intellect / imagination total score | 948 | 23.84 | 5.68 |
| NEO-FFI Neuroticism total value | 954 | 17.09 | 7.61 |
| NEO-FFI Extraversion total score | 943 | 26.97 | 5.92 |
| NEO-FFI Openness total score | 947 | 26.05 | 5.81 |
| NEO-FFI Agreeableness total score | 954 | 33.46 | 5.26 |
| NEO-FFI Conscientiousness total score | 947 | 34.66 | 5.97 |

| | | | |
|---|---|---|---|
| **PHYSICAL** | | | |
| Forced expiratory volume in 1 second | 1085 | 2.36 | 0.69 |
| Forced vital capacity | 1085 | 3.03 | 0.87 |
| Forced expiratory rate (FEV1/FVC ratio) | 1075 | 80.82 | 9.73 |
| Peak expiratory flow | 1085 | 350.72 | 133.94 |
| Grip strength (kg) best of 3 in right hand | 1085 | 28.96 | 10.14 |
| Grip strength (kg) best of 3 in left hand | 1084 | 27.09 | 10.1 |
| Height (cm) | 1090 | 166.42 | 8.93 |
| Weight in (kg) | 1089 | 77.12 | 14.26 |
| Body Mass Index (kg/m2) | 1089 | 27.78 | 4.36 |
| 6 metre walk time in seconds | 1085 | 3.86 | 1.16 |
| Demi-span (cm) | 1088 | 77.79 | 4.84 |
| Head circumference (cm) | 1089 | 57.03 | 2.05 |
| Townsend's Disability Scale score - Activities of daily living | 1089 | 1 | 1.97 |
| Telomere length | 1070 | 4200.51 | 559.67 |
| Cancer or tumour (yes/no) | 134/957 | - | - |
| Thyroid disorder (yes/no) | 99/991 | - | - |
| Parkinson's Disease (yes/no) | 5/1086 | - | - |
| Arthritis (yes/no) | 477/613 | - | - |
| Allergies (yes/no) | 333/758 | - | - |
| Medical history of gout or allopurinol medication (yes/no) | 38/1038 | - | - |
| Diagnosis of diabetes (yes/no) | 91/1000 | - | - |
| Leg pain when walking or in bed at night (yes/no) | 414/676 | - | - |
| **LIFESTYLE** | | | |
| Smoking category<br>- current<br>- ex<br>- never | <br>125<br>465<br>501 | - | - |
| Units of alcohol consumer per week | 1091 | 10.52 | 14.19 |
| Number of days a month exercise | 956 | 7.68 | 8.12 |
| Level of physical activity<br>- household chores<br>- walking etc 1-2 times a week<br>- walking etc several times a week<br>- exercise 1-2 times a week<br>- exercise several times a week<br>- keep-fit/heavy exercise/sport several times a week | <br>87<br>174<br>495<br>100<br>68<br>30 | - | - |
| Energy in kilo calories daily | 928 | 1912.7 | 650.5 |
| SIMD Deprivation Index | 1091 | 4564.74 | 1907.33 |
| Deprivation group (8 groups from SIMD) | 1083 | 6.25 | 2.1 |
| **LIFE-HISTORY** | | | |
| Age-11 IQ (age-11 Moray House Test score corrected for age in days, then converted to IQ) | 1028 | 100 | 14.99 |
| Number of years of full-time education | 1091 | 10.74 | 1.13 |
| Childhood social circumstances | 1082 | -0.0008 | 2.4 |
| Adult occupational social class | | - | - |

| | |
|---|---|
| - Professional<br>- Managerial/Technical<br>- Skilled (non-manual)<br>- Skilled (manual)<br>- Partly skilled<br>- Unskilled | 190<br>402<br>246<br>188<br>38<br>6 |
