## Supplementary file 2 for "Childhood intelligence attenuates the association between biological ageing and health outcomes in later"

**Supplementary file 2**. Data collection protocols for all phenotypes.

1) Blood

Blood samples were taken to assess:

- **$HbA_{1c}$** (%total)
- **Fibrinogen** (g/L)
- **C-Reactive Protein** (mg/L)
- **Triglycerides** (mmol/L)
- **Total serum cholesterol** (mmol/L)
- **High density lipoprotein cholesterol** (HDL; mmol/L)
- **HDL ratio** (HDL/total cholesterol)
- **Red cell count** (L)
- **White cell count** (L)
- **Neutrophils** (L)
- **Basophils** (L)
- **Monocytes** (L)
- **Lymphocytes** (L)
- **Eosinophils** (L)
- **Red cell folate** (ug/L)
- **Creatinine** (umol/L)
- **Total T3** (nmol/L)
- **Free T4** (pmol/L)
- **Thyroid-stimulating hormone** (mU/L)
- **Albumin** (g/L)
- **Calcium** (mmol/L)
- **Sodium** (mmol/L)
- **Vitamin B12** (ng/L)
- **Serum folate** (ug/L)
- **PT ratio**
- **APTT ratio**
- **Urea** (mmol/L)
- **Mean cell volume** (fL)

- **Platelet count** (L)
- **Prothrombin time** (secs)
- **Activated Partial Thromboplastin Time** (secs)
- **Potassium** (mmol/L)
- **Haemoglobin** (g/L)
- **Haematocrit** (ratio)

2) Cardiovascular

- **High blood pressure** – self reported (yes/no)
- **High cholesterol** - self reported (yes/no)
- **History of cardiovascular disease** - self reported (yes/no)
- **Problems with blood circulation** - self reported (yes/no)
- **History of stroke** - self reported (yes/no)
- **Family history of heart disease, stroke or problems with blood vessels** - self reported (yes/no)
- **Sitting and standing systolic and diastolic blood pressure** was measured using an Omron 705IT monitor (Milton Keynes, UK). Blood pressure scores were adjusted for individuals taking antihypertensive medications. The mean of the three measurements was included in analysis.

3) Cognitive

- **Mini-mental state examination (MMSE)** is a 30-point questionnaire that is typically used to measure cognitive impairment and screen for dementia (1). The test includes simple questions and covers areas of orientation to time, orientation to place, registration, attention and calculation, recall, language, repetition and complex commands. Scores indicate severe (≤12 point), moderate (13-19 points) or mild (20-24 points). Scores of ≥25 points is indicative of normal cognition.
- Six subtests of the Wechsler Adult Intelligence Scale III (2) were included:
  - **Digit Symbol Coding** was used to assess speed of information processing. This involves entering a symbol according to a number-symbol code, completing as many as possible in two minutes

- **Symbol Search** was also used to assess speed of information processing. This requires examination of a row of symbols to see if it contains one of a pair of target symbols, completing as many items as possible in the allotted time.
- **Matrix Reasoning** was used to assess non-verbal reasoning. Participants are asked to examine patterns displayed in a matrix and identify the missing item based on this rule.
**- Block Design** was used to assess constructional ability. This test requires participants to reconstruct specific designs from blocks with a maximum of two minutes per design.
**- Letter-Number Sequencing** was used to assess working memory. The test involves listening to increasingly long strings of numbers and letters and repeating these back in numerical and alphabetical order.
**- Backward Digit Span** was also used to assess working memory. Participants are required to listen to increasingly long strings of numbers and repeat them backwards.

- Tests of prior (or crystallised) cognitive ability included the **Wechsler Test of Adult Reading** (3) and the **National Adult Reading Test** (4). These tests comprise 50 written words designed to test the participant's vocabulary. The words typically have irregular spellings which do not have regular pronunciation rules.
- Three items from the Wechsler Memory Scale-III (5) were included:
  - **Logical memory** I and II (total score) – These tests assess immediate and delayed verbal declarative memory respectively. Logical memory I involves the immediate recollection of a story with 25 elements that is read aloud. Two stories are read with recall after each. The second story is read twice and participants are informed that they will be asked about the stories again later. Logical memory II involves recollecting as much as possible from the two stories read in the initial test.
  - **Spatial span** is a test of non-verbal, spatial learning and memory. The participant watches a tester touching the top of a number of blocks in a spatial array with the aim of touching the same blocks in the same order. The task is repeated with the aim of touching the blocks in the reverse order.
  - **Verbal paired associates** (total score) tests verbal learning and memory. Testers read a list of pairs of words in which some pairs have no obvious connection. They are then read the first of each pair and asked to recall the other. There are 8 word pairs and the task is repeated in different orders four times. Following a delay and without the pairs of words being read again the task is repeated.
- **Simple and four choice reaction time** (mean) – was used to assess speed of simple information processing. The tasks were administered using a stand-alone shallow

rectangular box constructed for the UK Health and Lifestyle Survey. This was described in detail previously (6). Briefly, there are five response keys numbered (from left to right) 1, 2, 0, 3, 4. In the simple reaction time test there are 8 practice trials and 20 test trials. The participant rests the second finger of the preferred hand on the 0 key. After a zero appears on the LCD screen the participant presses the key as fast as possible. The mean of the 20 simple reaction time trials are calculated. The four-choice reaction time test has 8 practice trials and 40 test trials. The participant rests the second and third fingers of the left and right hands on, respectively, the keys marked 1, 2, 3, 4. After a number appears on the LCD screen the participant presses the appropriate key as quickly as possible. Separate means are computed for correct and incorrect trials.

- **Inspection time** - is a two-alternative, forced choice, backward masking, visual discrimination task. It was used to assess speed of elementary visual processing. The inspection time task was replicated as closely as possible from a previous study (7), but with a longer instruction period and more practice trials. The participants were required to indicate which of two parallel, vertical lines of markedly different lengths was longer. The inspection time test was constructed, run, and analysed using E-Prime (Psychology Software Tools, Pittsburgh, PA). The stimulus lines were 5cm for the longer line and 2.5cm for the shorter line. They were joined at the top with a 2.5cm crossbar. Ten trials were presented at each of 15 durations (rounded to the nearest millisecond): 6, 12, 19, 25, 31, 37, 44, 50, 62, 75, 87, 100, 125, 150, and 200. Participants indicated the position of the longer line by pressing 1 or 2 on the number pad of a computer keyboard. The correctness of each response was noted.
- **Verbal fluency** was used to assess executive function. Participants are asked to name as many words as possible beginning with the letters C, F and L with one minute for each letter. No proper names are allowed and any words that are repeated only count once.

4) Life-history

- **Age 11 IQ** – was calculated from participants' Moray House Test score, corrected for age in days at the time of testing, and then converted to an IQ score. The Moray House Test was the test taken by most of the LBC1936 participants aged 11 as part of the Scottish Mental Survey 1947. The participants re-sat the test at Wave 1 using the same instructions and with the same time limit. The test has a variety of items including: following directions (14 items), same-opposites (11), word classification (10), analogies (8), practical items (6), reasoning (5), proverbs (4), arithmetic (4), spatial items (4), mixed sentences (3), cypher decoding (2), and other items (4).

- **Number of years of full-time education**
- **Occupational social class** was indexed according to the Classification of Occupations system (Her Majesty's Stationary Office (HMSO) code) which consists of 6 classes ranging from professional to unskilled (8). Married women's social class was assigned as their spouse's if higher than their own.
- **Age-11 social circumstances** – this was a measure of childhood deprivation and defined as the sum or the Z-scores of the number of rooms in the house divided by the number of people living in the house; whether the toilet was indoors or outdoors; and the number of people who shared the toilet at age 11.

5) Lifestyle

- Smoking category (current, ex, never)
- Units of alcohol consumed per week
- Level of physical activity – this was assessed on a 6-point scale, coded as follows:

  1) Household chores
  2) Walking etc. 1–2 times a week
  3) Walking etc. several times a week
  4) Exercise 1–2 times a week
  5) Exercise several times a week
  6) Keep-fit/heavy exercise/sport several times a week

- **Number of days a month of exercise** – number of days per month in which participants were active for 20 minutes or more in any vigorous sport or exercise
- **Energy in kilo calories daily** (derived from the Food Frequency Questionnaire)
- **Social deprivation** was measured using the Scottish Index of Multiple Deprivation (SIMD; 19). The SIMD ranks geographical areas in Scotland based on current income, employment, health, education, skills and training, geographic access to services, housing and crime. The SIMD provides a standardised measure of relative deprivation throughout Scotland.

6) Personality and mood

- **Hospital Anxiety and Depression Scale (HADS)** was used to measure recent mood state (9). The HADS is a fourteen item scale with 7 items relating to anxiety (A) and 7 to depression (D). The scale has a maximum score of 21, with probable anxiety or depression at scores of 11 or over.
- **The International Personality Item Pool (IPIP)** questionnaire (50-item) was used to assess personality traits (10). The questionnaire has 10 items each for 5 personality factors: Extraversion (E), Agreeableness (A), Conscientiousness (C), Emotional stability (ES) and Intellect (I). Participants were required to rate how well they believed each statement described them on a 5-point scale (from very inaccurate to very accurate).
- **The NEO Five Factor Inventory (NEO-FFI)** (11) is a 60-item inventory comprised of 12 items each for 5 factors: Neuroticism (N), Extraversion (E), Openness (O), Agreeableness (A) and Conscientiousness (C). Participants are required to mark each item as to how well it described them on a 5-point scale (from strongly disagree to strongly agree).

7) Physical

- **Height** (cm)
- **Weight** (kg)
- **BMI** (kg/m2)
- **Time taken to walk six metres** (sec)
- **Grip strength** in both right and left hands (kg; measured using a North Coast Hydraulic Hand Dynamometer, JAMAR)
- **Forced expiratory volume in 1 second (FEV1), forced vital capacity (FVC), forced expiratory ratio (FER (FEV1/FVC)), peak expiratory flow (PEF)** were all measured using a Micro Medical Spirometer, each the best of three.
- **Demispan** (cm)
- **Head circumference** (cm)
- **Townsend's Disability Scale - Activities of Daily Living (ADL)** is a 9-item index of activities that assess physical ability in social terms (12). Participants were given a score of 0 (no difficulty completing this activity) to 2 (not able to complete this activity) for each activity, and thus higher scores represent more functional disability.
- **Telomere length** - measured using a quantitative real-time polymerase chain reaction (performed on a 7900HT Fast Real Time PCR machine (Applied Biosystems; Pleasonton, CA,

USA)) assay. Plate-to-plate variation was corrected by running four internal control DNA samples on each plate.

- **Cancer or tumour** - self reported (yes/no)
- **Thyroid disorder** - self reported (yes/no)
- **Parkinson's Disease** - self reported (yes/no)
- **Arthritis** - self reported (yes/no)
- **Allergies** - self reported (yes/no)
- **Medical history of gout or allopurinol medication** - self reported (yes/no)
- **Diagnosis of diabetes** - self reported (yes/no)
- **Leg pain** when walking or in bed at night - self reported (yes/no)

*References*


1. Folstein MF, Folstein SE, McHugh PR. "Mini-mental state". A practical method for grading the cognitive state of patients for the clinician. Journal of psychiatric research. 1975;12(3):189-98.
2. Wechsler D. WAIS-IIIUK administration and scoring manual. Psychological Corporation. 1998.
3. Wechsler D. Wechsler Test of Adult Reading: WTAR. Psychological Corporation. 2001.
4. Nelson HE, Willison JR. National Adult Reading Test (NART) Test Manual (Part II) Windsor, UK. NFER-Nelson. 1991.
5. Wechsler D. WMS-IIIUK administration and scoring manual. Psychological Corporation. 1998.
6. Deary IJ, Der G, Ford G. Reaction times and intelligence differences: A population-based cohort study. Intelligence. 2001;29(5):389-99.
7. Deary IJ, Simonotto E, Meyer M, Marshall A, Marshall I, Goddard N, et al. The functional anatomy of inspection time: an event-related fMRI study. NeuroImage. 2004;22(4):1466-79.
8. OPCS. Classification of occupations and coding index. Her Majesty's Stationery Office London; 1980.
9. Zigmond AS, Snaith RP. The hospital anxiety and depression scale. Acta psychiatrica Scandinavica. 1983;67(6):361-70.
10. Available from: http://ipip.ori.org/.
11. Costa PT, McCrae RR. NEO PI-R Professional Manual. Psychological Assessment Resources. 1992.
12. Townsend P. Poverty in the United Kingdom: a survey of household resources and standards of living. Penguin. 1979.
