## Supplementary file 3 for "Childhood intelligence attenuates the association between biological ageing and health outcomes in later"

**Supplementary file 3.** Associations between DNAm PhenoAgeAccel and all phenotypes.

**Figure 1.** Associations between DNAm PhenoAgeAccel and all the blood-based variables at Wave 1. Standardised model β coefficients (effect sizes) are presented along the x-axis. Phenotypes are presented along the y-axis. Error bars show the 95% confidence interval. Phenotypes marked with an asterisk (*) are significantly associated with DNAm PhenoAgeAccel at an FDR-corrected p-value of <0.05.

*Abbreviations*: HDL: high-density lipoprotein; APTT: activated partial thromboplastin time; TSH: thyroid stimulating hormone; CRP: C-reactive protein

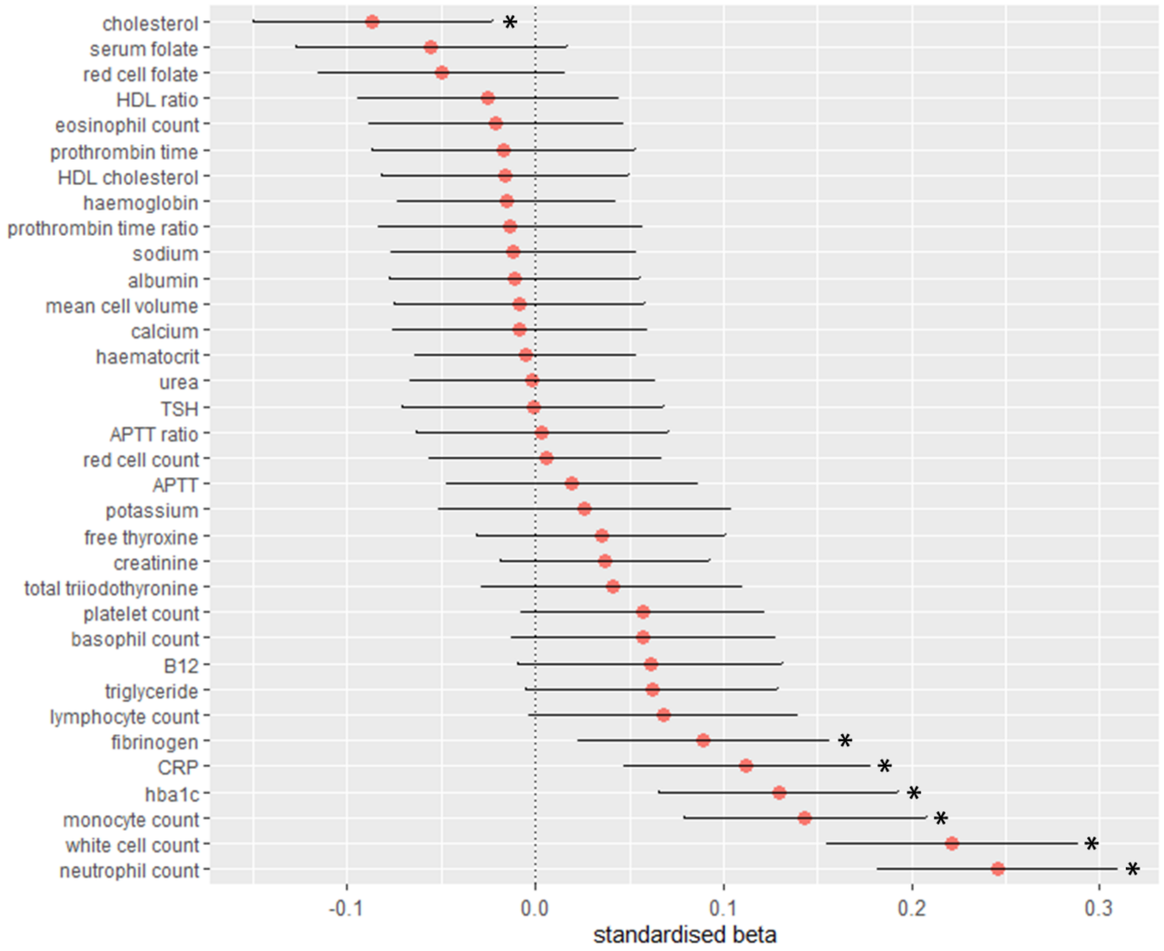

**Figure 2**. Associations between DNAm PhenoAgeAccel and all cardiovascular variables at Wave 1. Standardised model β coefficients (for continuous variables) or log odds (for binary variables) are presented along the x-axis. Phenotypes are presented along the y-axis. Error bars show the 95% confidence interval.

*Abbreviations:* family history: family history of heart disease, stroke, or problems with blood vessels; DBP: diastolic blood pressure; CVD: cardiovascular disease; SBP: systolic blood pressure.

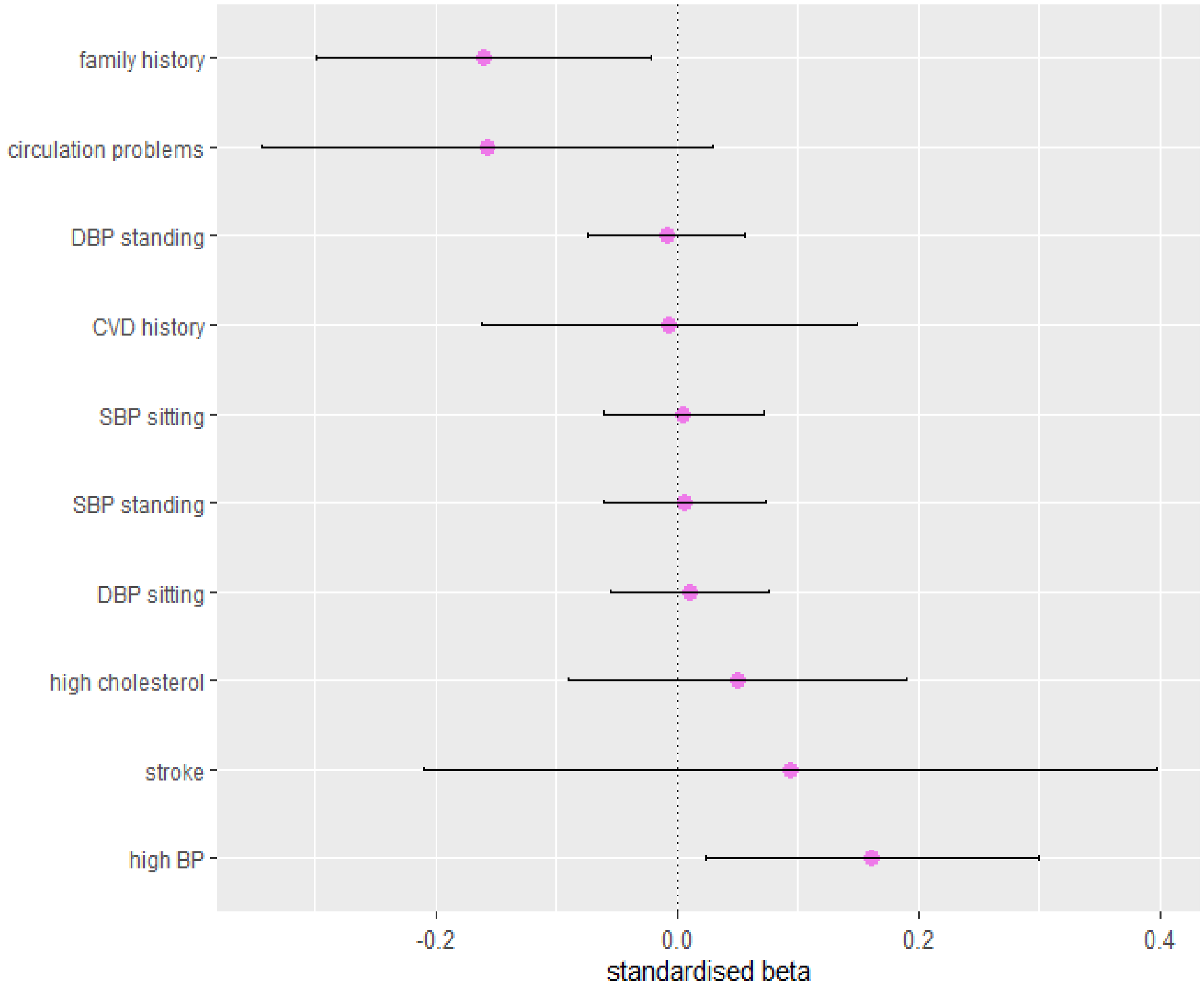

**Figure 3**. Associations between DNAm PhenoAgeAccel and all cognitive variables at Wave 1. Standardised model β coefficients are presented along the x-axis. Phenotypes are presented along the y-axis. Error bars show the 95% confidence interval. Phenotypes marked with an asterisk (*) are significantly associated with DNAm PhenoAgeAccel at an FDR-corrected p-value of <0.05.

*Abbreviations:* MMSE: mini-mental state examination; VPA: verbal paired associates; WTAR: Wechsler Test of Adult Reading; RT: reaction time.

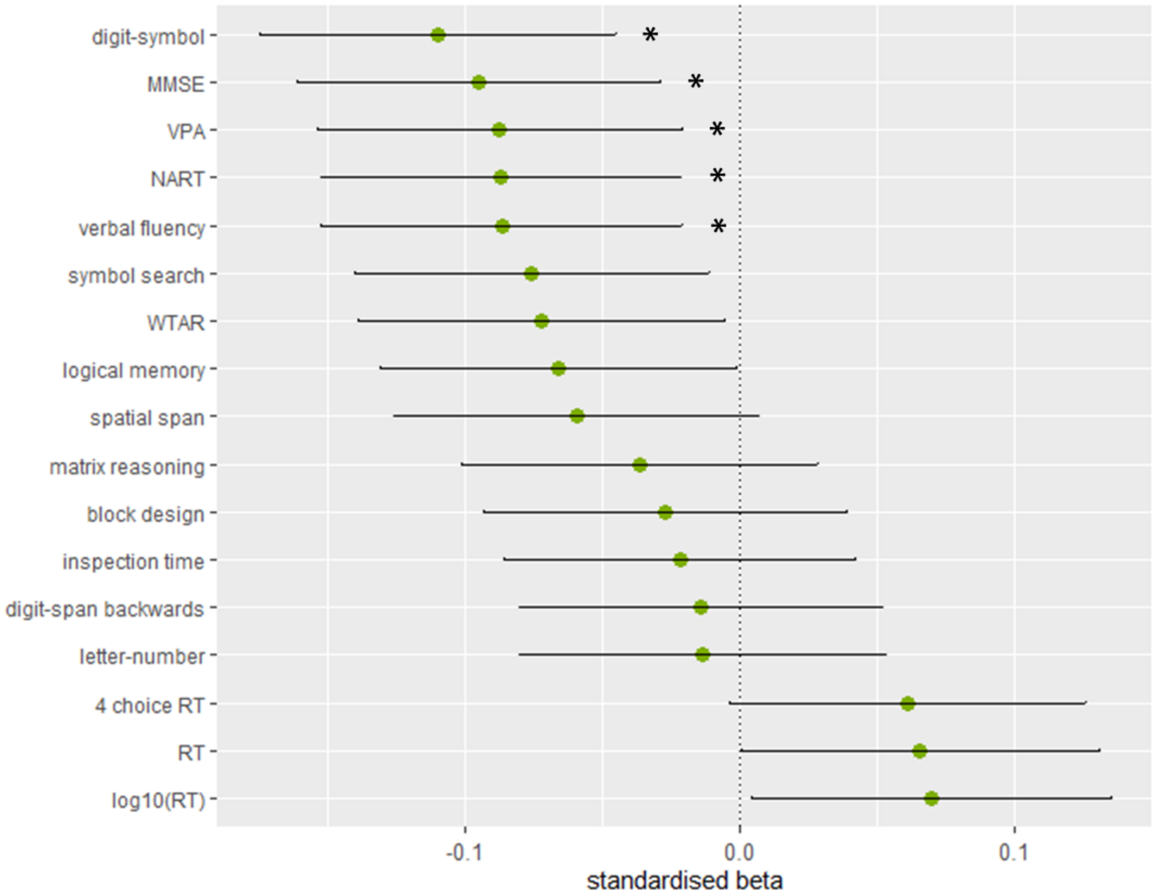

**Figure 4.** Associations between DNAm PhenoAgeAccel and all personality and mood phenotypes at Wave 1. Standardised model β coefficients are presented along the x-axis. Phenotypes are presented along the y-axis. Error bars show the 95% confidence interval.

*Abbreviations:* NEO: Neuroticism-Extraversion-Openness Five-Factor Inventory; IPIP: International Personality Item Pool; HADS: Hospital Anxiety and Depression Scale.

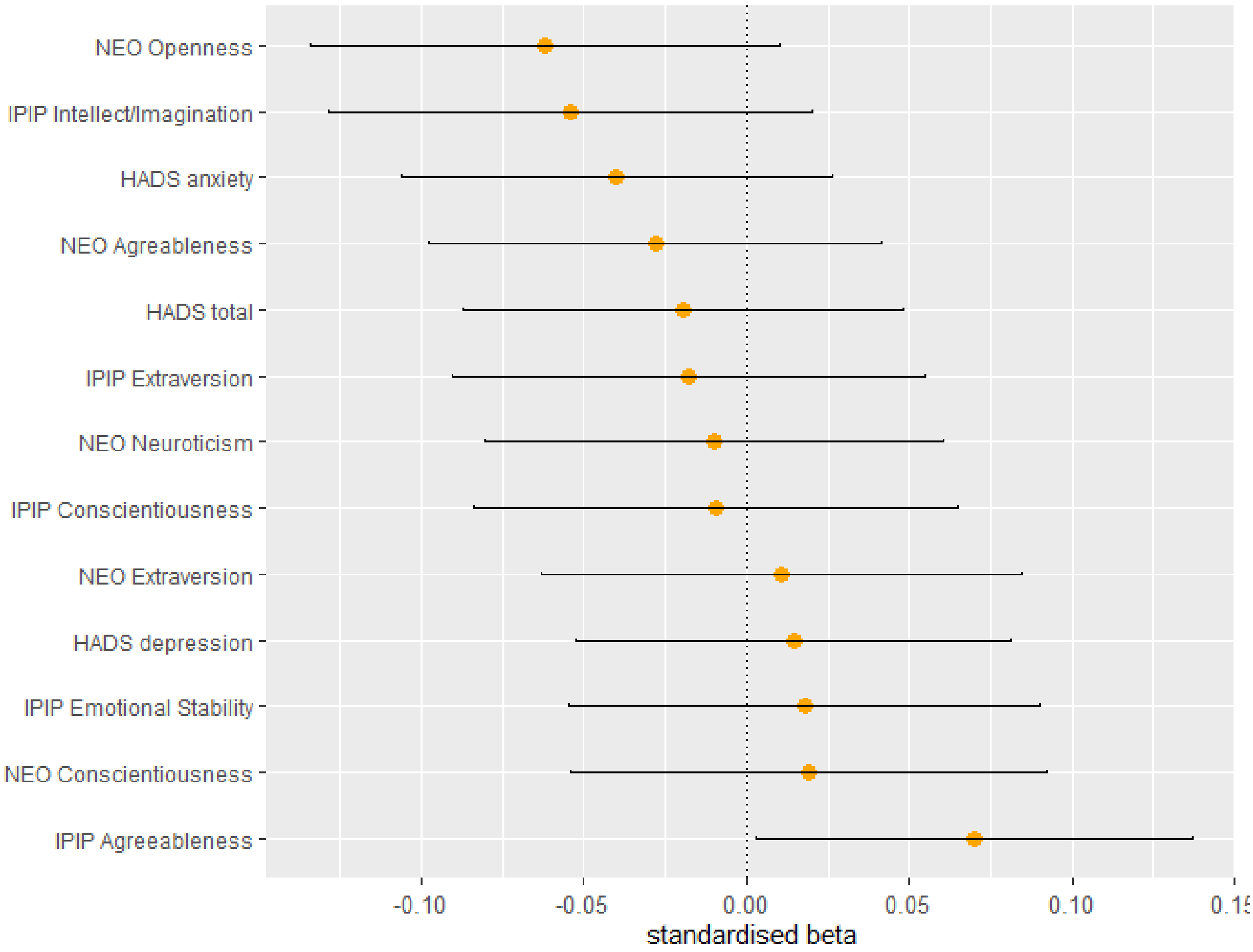

**Figure 5**. Associations between DNAm PhenoAgeAccel and all physical phenotypes at Wave 1. Standardised model β coefficients (for continuous variables) or log odds (for binary variables) are presented along the x-axis. Phenotypes are presented along the y-axis. Error bars show the 95% confidence interval. Phenotypes marked with an asterisk (*) are significantly associated with DNAm PhenoAgeAccel at an FDR corrected p-value of <0.05.

*Abbreviations:* FEV1: forced expiratory volume in 1 second; FVC: forced vital capacity; PEF: peak expiratory flow; FER: forced expiratory volume; BMI: body mass index; ADL: activities of daily living (Townsend Disability Scale).

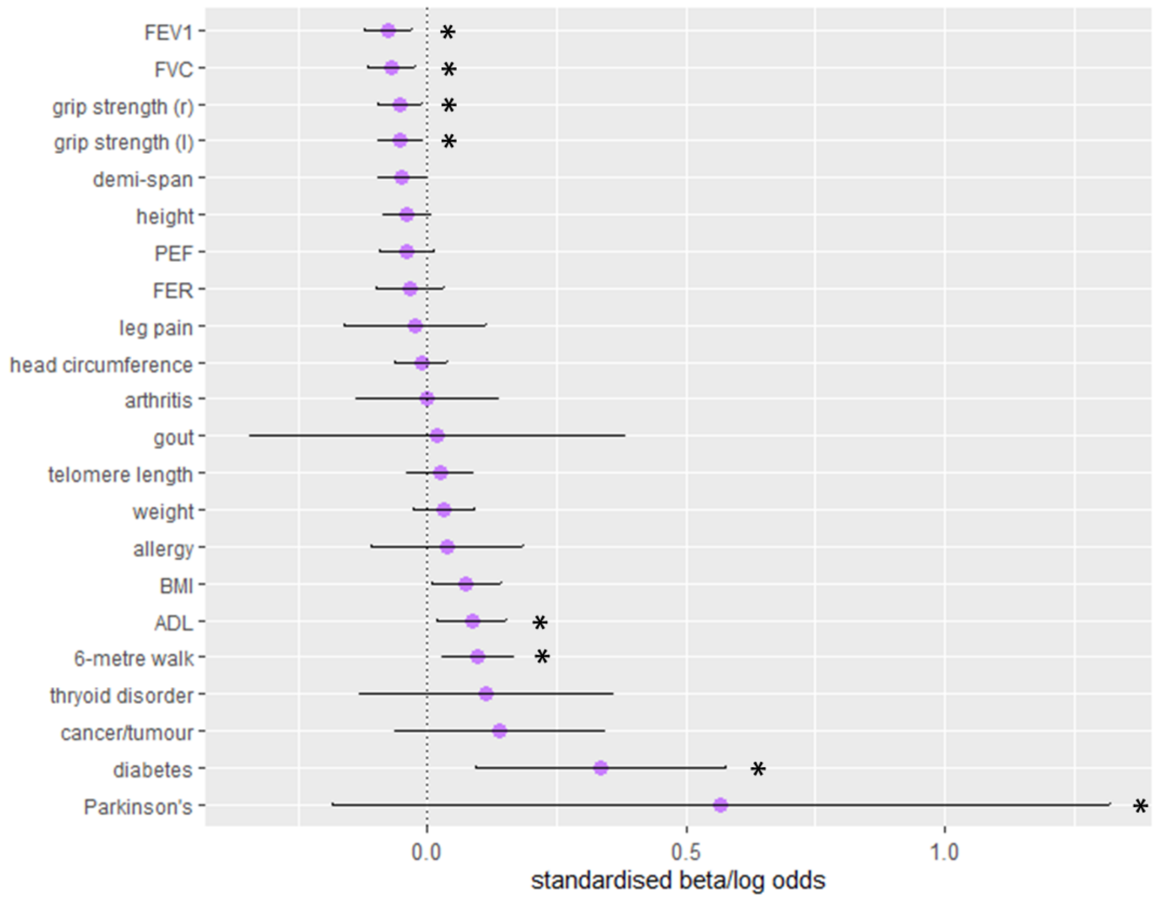

**Figure 6**. Associations between DNAm PhenoAgeAccel and all lifestyle phenotypes at Wave 1. Standardised model β coefficients (for continuous variables) or log odds (for binary or ordinal variables) are presented along the x-axis. Phenotypes are presented along the y-axis. Error bars show the 95% confidence interval. Phenotypes marked with an asterisk (*) are significantly associated with DNAm PhenoAgeAccel at an FDR-corrected p-value of <0.05. The reference level for smoking category was never smokers.

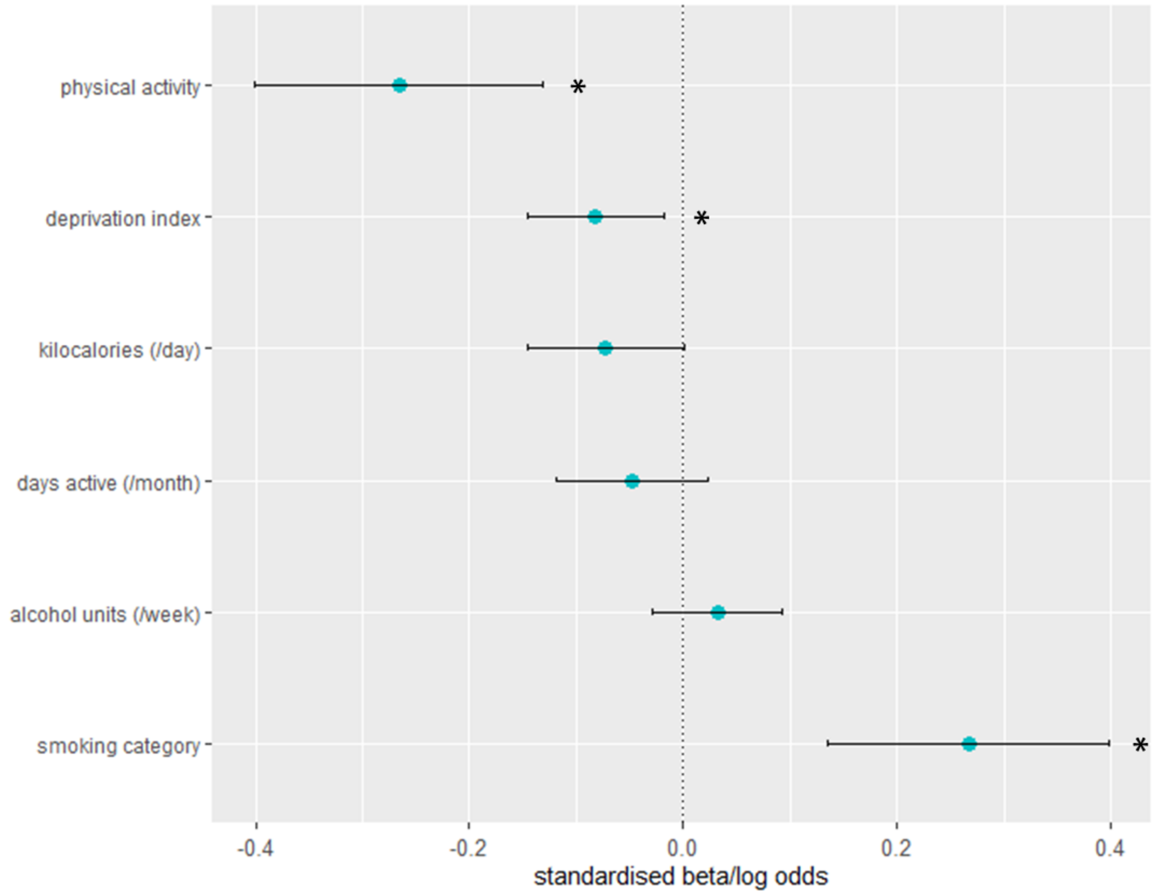

**Figure 7**. Associations between DNAm PhenoAgeAccel and all life-history phenotypes at Wave 1. Standardised model β coefficients (for continuous variables) or log odds (for binary or ordinal variables) are presented along the x-axis. Phenotypes are presented along the y-axis. Error bars show the 95% confidence interval. Phenotypes marked with an asterisk (*) are significantly associated with DNAm PhenoAgeAccel at an FDR corrected p-value of <0.05.

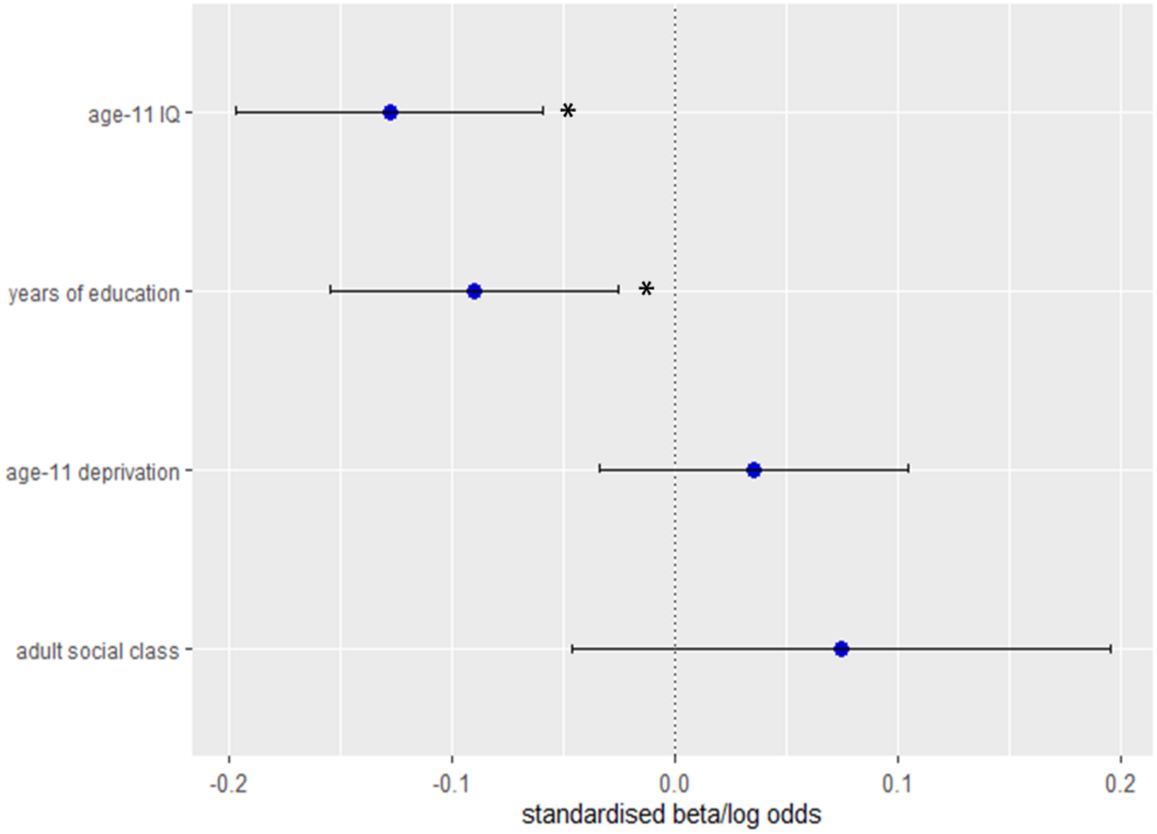
