## Supplementary file 4 for "Childhood intelligence attenuates the association between biological ageing and health outcomes in later"

**Supplementary file 4**. Full PheWAS results.

*Abbreviations*: FDR: False Discovery Rate; HADS: Hospital Anxiety and Depression Scale WMS: Wechsler Memory Scale; WAIS: Wechsler Adult Intelligence Scale; IPIP: International Personality Item Pool; NEO-FFI: Neuroticism-Extraversion-Openness Five-Factor Inventory; SIMD: Scottish Index of Multiple Deprivation.

| **Phenotype** | **Standardised β/ log odds** | **Standard error** | **Raw p** | **FDR-corrected p** |
|---|---|---|---|---|
| **BLOOD** | | | | |
| Haemoglobin | -0.015 | 0.030 | 0.607 | 0.904 |
| Red cell count | 0.005 | 0.032 | 0.865 | 0.949 |
| Haematocrit | -0.006 | 0.030 | 0.852 | 0.949 |
| Mean cell volume | -0.009 | 0.034 | 0.800 | 0.949 |
| White cell count | 0.222 | 0.034 | **<0.001** | **<0.001** |
| Neutrophil count | 0.245 | 0.032 | **<0.001** | **<0.001** |
| Lymphocyte count | 0.068 | 0.036 | 0.063 | 0.262 |
| Monocyte count | 0.143 | 0.037 | **<0.001** | **<0.001** |
| Eosinophil count | -0.021 | 0.034 | 0.537 | 0.904 |
| Basophil count | 0.057 | 0.036 | 0.108 | 0.306 |
| Platelet count | 0.057 | 0.033 | 0.085 | 0.275 |
| Prothrombin time | -0.017 | 0.036 | 0.638 | 0.904 |
| Activated partial thromboplastin time | 0.019 | 0.034 | 0.568 | 0.904 |
| Fibrinogen | 0.089 | 0.034 | **0.009** | **0.043** |
| Vitamin B12 | 0.061 | 0.036 | 0.089 | 0.275 |
| Serum folate | -0.055 | 0.037 | 0.134 | 0.337 |
| Red cell folate | -0.050 | 0.034 | 0.139 | 0.337 |
| Prothrombin time ratio | -0.013 | 0.036 | 0.707 | 0.936 |
| Activated partial thromboplastin time ratio | 0.004 | 0.034 | 0.916 | 0.971 |
| Urea | -0.002 | 0.033 | 0.958 | 0.971 |
| Creatinine | 0.037 | 0.028 | 0.192 | 0.437 |
| Sodium | -0.012 | 0.033 | 0.716 | 0.936 |
| Potassium | 0.026 | 0.040 | 0.514 | 0.904 |
| Albumin | -0.011 | 0.034 | 0.750 | 0.945 |
| Calcium | -0.008 | 0.034 | 0.813 | 0.949 |
| Cholesterol | -0.086 | 0.032 | **0.008** | **0.043** |
| Triglyceride | 0.062 | 0.034 | 0.069 | 0.262 |
| High-density lipoprotein cholesterol | -0.016 | 0.034 | 0.626 | 0.904 |
| Cholesterol: HDLC Ratio | -0.026 | 0.035 | 0.467 | 0.882 |
| $HbA_{1C}$ | 0.129 | 0.032 | **<0.001** | **<0.001** |

| | | | | |
|---|---|---|---|---|
| C-reactive protein | 0.112 | 0.033 | **0.001** | **0.006** |
| Thyroid stimulating hormone | -0.001 | 0.035 | 0.971 | 0.971 |
| Free thyroxine | 0.035 | 0.034 | 0.298 | 0.596 |
| Total triiodothyronine | 0.041 | 0.036 | 0.250 | 0.530 |
| **CARDIOVASCULAR** | | | | |
| Mean diastolic blood pressure - sitting | 0.011 | 0.034 | 0.734 | 0.940 |
| Mean diastolic blood pressure - standing | -0.009 | 0.033 | 0.792 | 0.940 |
| Mean systolic blood pressure - sitting | 0.006 | 0.034 | 0.868 | 0.940 |
| Mean systolic blood pressure - standing | 0.007 | 0.034 | 0.848 | 0.940 |
| High blood pressure | 0.162 | 0.070 | **0.021** | 0.121 |
| High cholesterol | 0.050 | 0.072 | 0.484 | 0.940 |
| Cardiovascular disease history | -0.006 | 0.079 | 0.940 | 0.940 |
| Problems with blood circulation | -0.157 | 0.095 | 0.100 | 0.333 |
| History of stroke | 0.094 | 0.155 | 0.544 | 0.940 |
| Family history of heart disease, stroke or problems with blood vessels | -0.160 | 0.071 | **0.024** | 0.121 |
| **COGNITIVE** | | | | |
| MMSE | -0.095 | 0.034 | **0.005** | **0.034** |
| WMS III - Logical memory | -0.066 | 0.033 | **0.047** | 0.083 |
| WMS III - Spatial span total | -0.059 | 0.034 | 0.079 | 0.112 |
| WMS III - Verbal paired associates | -0.087 | 0.034 | **0.009** | **0.034** |
| WAIS III - Symbol search | -0.076 | 0.033 | **0.021** | 0.060 |
| WAIS III - Digit-symbol coding | -0.110 | 0.033 | **<0.001** | **0.015** |
| Simple reaction time mean | 0.065 | 0.033 | **0.049** | 0.083 |
| Log10 simple reaction time | 0.070 | 0.033 | **0.037** | 0.079 |
| Four choice reaction time | 0.061 | 0.033 | 0.065 | 0.101 |
| Inspection time total correct responses | -0.022 | 0.033 | 0.504 | 0.571 |
| WAIS III - Matrix reasoning | -0.036 | 0.033 | 0.273 | 0.357 |

| | | | | |
|---|---|---|---|---|
| Verbal fluency | -0.087 | 0.033 | **0.009** | **0.034** |
| WAIS III - Letter-number sequencing | -0.013 | 0.034 | 0.692 | 0.692 |
| WAIS III - Digit span backwards | -0.014 | 0.034 | 0.677 | 0.692 |
| National Adult Reading Test | -0.087 | 0.033 | **0.009** | **0.034** |
| Wechsler Test of Adult Reading | -0.072 | 0.034 | **0.033** | 0.079 |
| WAIS III - Block design | -0.027 | 0.034 | 0.418 | 0.507 |
| **LIFESTYLE** | | | | |
| Units of alcohol/week | 0.032 | 0.031 | 0.307 | 0.307 |
| Number of days exercise/month | -0.048 | 0.036 | 0.190 | 0.228 |
| Energy in kilo calories/day | -0.072 | 0.038 | 0.054 | 0.082 |
| SIMD | -0.082 | 0.033 | **0.012** | **0.025** |
| Smoking category (current, ex, never) | 0.267 | 0.067 | **<0.001** | **<0.001** |
| Level of physical activity | -0.266 | 0.069 | **<0.001** | **<0.001** |
| **PERSONALITY AND MOOD** | | | | |
| HADS - Anxiety score | -0.040 | 0.034 | 0.236 | 0.767 |
| HADS - Depression score | 0.015 | 0.034 | 0.670 | 0.803 |
| HADS - Total score | -0.019 | 0.034 | 0.572 | 0.803 |
| IPIP Extraversion | -0.018 | 0.037 | 0.634 | 0.803 |
| IPIP Agreeableness | 0.070 | 0.034 | **0.042** | 0.543 |
| IPIP Conscientiousness | -0.010 | 0.038 | 0.803 | 0.803 |
| IPIP Emotional stability | 0.018 | 0.037 | 0.630 | 0.803 |
| IPIP Intellect / imagination | -0.054 | 0.038 | 0.154 | 0.667 |
| NEO-FFI Neuroticism | -0.010 | 0.036 | 0.783 | 0.803 |
| NEO-FFI Extraversion | 0.011 | 0.038 | 0.776 | 0.803 |
| NEO-FFI Openness | -0.062 | 0.037 | 0.094 | 0.611 |
| NEO-FFI Agreeableness | -0.028 | 0.036 | 0.430 | 0.803 |
| NEO-FFI Conscientiousness | 0.019 | 0.037 | 0.609 | 0.803 |
| **PHYSICAL** | | | | |
| Height | -0.041 | 0.023 | 0.073 | 0.161 |
| Weight | 0.032 | 0.030 | 0.280 | 0.441 |
| Body Mass Index | 0.075 | 0.034 | **0.027** | 0.075 |
| 6 metre walk time | 0.097 | 0.035 | **0.006** | **0.037** |
| Demi-span | -0.049 | 0.024 | **0.047** | 0.114 |
| Head circumference | -0.012 | 0.026 | 0.627 | 0.727 |

| | | | | |
|---|---|---|---|---|
| Townsend's Disability Scale score - Activities of daily living | 0.086 | 0.035 | **0.013** | **0.045** |
| Forced expiratory volume in 1 second | -0.108 | 0.025 | **<0.001** | **0.023** |
| Forced vital capacity | -0.089 | 0.023 | **0.002** | **0.023** |
| Forced expiratory rate (FEV1/FVC ratio) | -0.034 | 0.034 | 0.307 | 0.451 |
| Peak expiratory flow | -0.039 | 0.026 | 0.135 | 0.253 |
| Grip strength - right hand | -0.053 | 0.022 | **0.014** | **0.045** |
| Grip strength - left hand | -0.053 | 0.021 | **0.014** | **0.045** |
| Telomere length | 0.024 | 0.032 | 0.463 | 0.599 |
| Diagnosis of diabetes | 0.335 | 0.124 | **0.007** | **0.038** |
| Leg pain | -0.025 | 0.070 | 0.724 | 0.796 |
| Cancer or tumour | 0.139 | 0.104 | 0.178 | 0.302 |
| Thyroid disorder | 0.113 | 0.125 | 0.366 | 0.504 |
| Parkinson's Disease | 0.568 | 0.383 | 0.138 | 0.253 |
| Arthritis | -0.001 | 0.070 | 0.990 | 0.989 |
| Allergies | 0.038 | 0.075 | 0.616 | 0.726 |
| Medical history of gout or allopurinol medication | 0.018 | 0.185 | 0.922 | 0.966 |
| **LIFE-HISTORY** | | | | |
| Age-11 IQ | -0.128 | 0.035 | 0.001 | **0.001** |
| Years of education | 0.090 | 0.033 | 0.006 | **0.012** |
| Childhood social circumstances | 0.035 | 0.035 | 0.317 | 0.317 |
| Adult occupational social class | 0.074 | 0.062 | 0.227 | 0.302 |
