## Supplementary file 5 for "Childhood intelligence attenuates the association between biological ageing and health outcomes in later"

**Supplementary file 5**. Change in phenotypes over time (standard deviation per year).

| **Phenotype** | **Standardised β** | **Standard error** | **p** |
|---|---|---|---|
| *Grip strength (r)* | -0.035 | 0.002 | $<2x10^{-16}$ |
| *Grip strength (l)* | -0.031 | 0.002 | $<2x10^{-16}$ |
| *Forced expiratory volume (1s)* | -0.059 | 0.002 | $<2x10^{-16}$ |
| *Forced vital capacity* | -0.039 | 0.002 | $<2x10^{-16}$ |
| *Forced expiratory ratio* | -0.033 | 0.004 | $<2x10^{-16}$ |
| *Peak expiratory flow* | -0.053 | 0.003 | $<2x10^{-16}$ |
| *Digit span backwards* | -0.017 | 0.003 | $2x10^{-8}$ |
| *Symbol search* | -0.042 | 0.003 | $<2x10^{-16}$ |
| *Digit symbol coding* | -0.063 | 0.002 | $<2x10^{-16}$ |
| *Matrix reasoning* | -0.028 | 0.003 | $<2x10^{-16}$ |
| *Letter number sequencing* | -0.044 | 0.003 | $<2x10^{-16}$ |
| *Block design* | -0.044 | 0.003 | $<2x10^{-16}$ |
| *6m walk time (s)* | 0.099 | 0.003 | $<2x10^{-16}$ |
| *Telomere length* | -0.088 | 0.003 | $<2x10^{-16}$ |
